# Accounting for experimental noise reveals that mRNA levels, amplified by post-transcriptional processes, largely determine steady-state protein levels in yeast

**DOI:** 10.1101/009472

**Authors:** Gábor Csárdi, Alexander Franks, David S. Choi, Edoardo M. Airoldi, D. Allan Drummond

## Abstract

Cells respond to their environment by modulating protein levels through mRNA transcription and post-transcriptional control. Modest observed correlations between global steady-state mRNA and protein measurements have been interpreted as evidence that mRNA levels determine roughly 40% of the variation in protein levels, indicating dominant post-transcriptional effects. However, the techniques underlying these conclusions, such as correlation and regression, yield biased results when data are noisy, missing systematically, and collinear—properties of mRNA and protein measurements—which motivated us to revisit this subject. Noise-robust analyses of 24 studies of budding yeast reveal that mRNA levels explain more than 85% of the variation in steady-state protein levels. Protein levels are not proportional to mRNA levels, but rise much more rapidly. Regulation of translation suffices to explain this nonlinear effect, revealing post-transcriptional amplification of, rather than competition with, transcriptional signals. These results substantially revise widely credited models of protein-level regulation, and introduce multiple noise-aware approaches essential for proper analysis of many biological phenomena.

## Author Summary

Cells respond to their environment by making proteins using transcription and translation of mRNA. Modest observed correlations between global steady-state mRNA and protein measurements have been interpreted as evidence that mRNA levels determine roughly 40% of the variation in protein levels, indicating dominant post-transcriptional effects. However, the techniques underlying these conclusions, such as correlation and regression, yield biased results when data are noisy and contain missing values. Here we show that when methods that account for noise are used to analyze much of the same data, mRNA levels explain more than 85% of the variation in steady-state protein levels. Protein levels are not proportional to mRNA levels as commonly assumed, but rise much more rapidly. Regulation of translation achieves amplification of, rather than competition with, transcriptional signals. Our results suggest that for this set of conditions, mRNA sets protein-level regulation, and introduce multiple noise-aware approaches essential for proper analysis of many biological phenomena.

## Introduction

Cellular protein levels reflect the balance of mRNA levels, protein production by translation initiation and completion, and protein removal by degradation, secretion and dilution [1,2] (Figure 1A). A standard quantitative model for protein-level regulation [3,4] is

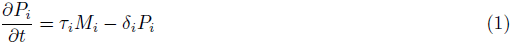

where *P*_*i*_ is the cellular protein level (molecules per cell) of gene *i, M*_*i*_ is the mRNA level, and *τ*_*i*_ and *δ*_*i*_ are the mRNA translation and net protein removal rates, respectively. According to this model, at steady-state, protein levels will be proportional to mRNA levels with proportionality constants of *τ*_*i*_/δ_*i*_:

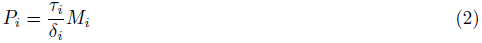

such that if rates of translation and removal did not vary by gene, and in the absence of experimental noise or other variation, steady-state mRNA and protein levels would correlate perfectly [1]. Consequently, the mRNA-protein correlation observed in global measurements of mRNA and protein levels has been intensely studied, and deviations from perfect correlation used to quantify the contribution of post-transcriptional processes to cellular protein levels [1,2,5–8].

**Figure 1.**
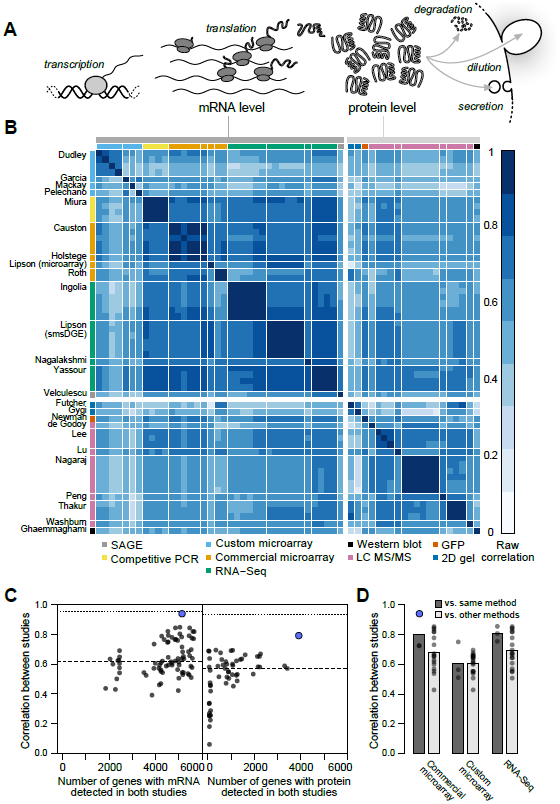
Measurements of steady-state mRNA and protein levels in budding yeast reveal wide variation in reproducibility and coverage. **A**, Steady-state protein levels reflect the balance of mRNA translation and protein removal. **B**, Raw correlations between measurements of mRNA and protein arranged by study (denoted by the first author) with the quantification method indicated. **C**, Measurements vary widely in reproducibility and coverage. Each point represents a pair of studies. Dots show between-study correlations (median shown by dashed line), a measure of reliability. Dotted line, median of within-study correlations. Blue dots show pairs of studies from the same research group. **D**, Correlations between studies sharing the same quantification method or different methods (dark and light gray bars, respectively), using mRNA datasets with ≥ 5000 genes (4,595 genes quantified by all datasets). For example, the second column from the left shows the 18 correlations between each of three commercial microarray studies and six studies using custom microarrays or RNA-Seq.

The consensus across these studies holds that, in a wide array of organisms, transcriptional regulation explains 30–50% of the variation in steady-state protein levels, leaving half or more to be explained by post-transcriptional regulatory processes [2,6,8–15]. Higher correlations are observed, generally for subsets of less than half the genome [1,8,16]. Low observed mRNA-protein correlations have motivated the search for alternate forms of regulation capable of accounting for the majority of protein-level variability [2,8,12]. In one proposal, mRNA levels serve mainly as an on-off switch for protein expression, imposing coarse control over protein levels which is then tuned by post-transcriptional mechanisms [8]. Recent studies have indeed uncovered wide between-gene variation in post-transcriptional features such as inferred translation rates [17] and protein degradation rates [2].

However, as frequently noted [1,6,8,9,18–20], noise in measurements can cause many of the observations attributed to post-transcriptional regulation. Here, noise encompasses variability due to cell-to-cell variation, growth conditions, sample preparation and other effects due to experimental design [21], and measurement biases and error [9,20]. Uncorrelated noise between mRNA and protein measurements will reduce the observed mRNA-protein correlation relative to the true value [22], while inflating the variation in measurements of translational efficiency and other post-transcriptional processes.

Most studies, particularly of protein levels, cover only a subset of known genes, due to factors such as signal-to-noise limitations, method biases, and continual revision of the coding-sequence annotations used to design and analyze assays. Limited and variable transcriptome and proteome coverage complicate analyses further, making it difficult to compare studies and to synthesize a holistic view of regulatory contributions. Missing data tends to reduce the precision of estimates, if data are missing at random (MAR). However, most quantification methods are biased toward detection of more abundant mRNAs and proteins [8]. Data which are not missing at random (NMAR) in this way have reduced variance or restricted range. Range restriction, in turn, tends to systematically attenuate (reduce in absolute magnitude toward zero) the observed correlations and regression coefficients relative to complete data [23,24]. That is, biased detection produces biased estimates of the mRNA-protein correlation, leading to incorrect conclusions about regulatory contributions [25].

In many comparisons of the roles of transcriptional and post-transcriptional regulation, protein levels are correlated with or regressed on various predictors (mRNA level and half-life, codon usage, amino-acid usage, *etc.*) to determine relative contributions to protein-level variation [1–3,13,17,20]. If mRNA levels are found to explain a certain percentage, say X, in protein levels, then the other predictors are asserted to explain no more than 100 – X percent of the variance [2,8,20,26]. A basic assumption of such analyses is that transcriptional and post-transcriptional regulation vary independently between genes. Several of the same studies report that high-expression genes show signs of more efficient translation [2, 3,17] (reviewed in [1]), raising concerns about the validity of this assumption.

A related assumption of these analyses, one encoded in the standard functional model above, is that mRNA and protein levels are proportionally or linearly related [1,4]; the slope of this line is the mean number of proteins per mRNA. More often, the data are plotted on a log-log scale, where linearity appears as a slope of 1. Consistent with this, ordinary least-squares linear regression shows that the slope is quite close to 1 for *E. coli* (0.96) and budding yeast (1.08) [16], and estimates of proteins per mRNA have been reported roughly constant across mRNA expression levels in a prominent study [27].

However, like correlations, slopes estimated by standard linear regression are biased downward by noise in mRNA level measurements, an effect called *regression dilution bias* [28] which affects any regression where the independent variable is measured with error. A frequently encountered case is that, given two measurements *X* and *Y*, the slope from regressing *Y* on *X* is not the inverse of the slope of regressing *X* on *Y* [29–31]; this is regression dilution bias at work. Consequently, linear regression cannot be used to estimate the functional relationship between mRNA and protein levels, raising the question of what the true functional relationship is. Use of nonparametric methods avoids assumptions of linearity [1], at the cost of destroying genuine information about the dynamic range of gene expression and its determinants.

Analytical solutions to many of these problems exist—notably, Spearman introduced a correction for noise-induced attenuation of correlation estimates more than a century ago [22]—yet have largely failed to find their way into the hands of groups carrying out gene-regulation experiments and analyses (with a few exceptions [14]). Some problems remain almost entirely unaddressed, such as providing accurate estimates of the functional relationship between variables measured many times with correlated noise yielding variably and systematically missing values.

Here, we develop and integrate approaches to address all of these challenges, with the aim of providing more comprehensive and rigorous estimates of the relationship between mRNA and protein levels than have previously been possible. To do so, we take advantage of the rapid, continual progress made in global measurement of mRNA and protein levels by multiple methods [5,16,27,32–38]. All of these methods were first employed at the genome scale in studies profiling gene expression during log-phase growth of budding yeast in rich medium, a *de facto* standard. These studies often compare results against previous studies, evaluating agreement, precision, coverage, and dynamic range while pointing out relative advantages of each approach (*e.g.* [16,17,27,34,35,37]).

Our efforts to synthesize these data into a coherent whole are grounded in the stance that all these works constitute measurements of the same underlying quantities—average mRNA and protein levels in a large cell population prepared under narrowly defined conditions—whether or not such measurements were the study goal. Systematic differences between approaches due to experimental choices will introduce variation which may not be distinguishable from simple inaccuracy in measurement. We treat this variation as experimental noise without prejudice. Distinctions between biological variability, measurement error, method bias, and other sources of noise are of course important, particularly in deciding how to control or manage noise. These distinctions may also depend on one’s perspective. For example, unintentional differences in growth conditions may lead two groups following the same protocol to make measurements on samples which inevitably are, in truth, biologically different, such that error-free measurement would reveal differences in mRNA and protein levels. In one sense, these differences reflect biological variability; in an equally valid sense, they represent experimental noise. Similarly, intentional protocol differences that are not meant to alter measurement accuracy (such as use of new methods intended to make measurements more precise), yet carry known and unknown biases, may also introduce noise. Here, we take an empirical approach to noise which does not involve divining intent. Versions of this approach are taken, often implicitly, by the many previous analyses that integrate experiments from multiple groups [7,8,16,17,27].

Our results reveal that, once noise is accounted for, mRNA and protein levels correlate much more strongly under these experimental conditions than previously appreciated, with a correlation coefficient of *r* = 0.93. We find that protein levels are not proportional to mRNA levels, but instead are more steeply related, an effect we show is consistent with measurements of translational activity. Transcriptional and post-transcriptional regulation act in a concerted, non-independent manner to set protein levels, inconsistent with common attempts to divvy up and assign protein-level variance to each mechanism. As a byproduct, we generate what by several measures is the most complete and accurate quantitative transcriptome and proteome available, in average molecules per haploid cell, for this widely studied organism under these well-studied conditions. Finally, we highlight and introduce methods for analyzing correlations and functional relationships between measured data which may be used broadly.

## Results

### Correlations and coverage range widely across datasets

We collected 38 measurements of mRNA levels and 20 measurements of protein levels from 13 and 11 separate studies respectively, each of haploid *S. cerevisiae* growing exponentially in shaken liquid rich medium with 2% glucose between 22°C and 30°C (Table 1). As described in the Introduction, we assume, for modeling purposes, that each replicate in each experiment constitutes a measurement of the true pergene mean mRNA and protein levels under these narrowly defined conditions. These data cover varying amounts of the genome and display a wide range of correlations between studies (Figure 1B, Pearson correlations on log-transformed values with zeros and missing values omitted).

**Table 1.** Measurements of mRNA (above the midline) and protein (below the midline) analyzed using structured covariance modeling.

| Study identifier [reference] | Technology (replicates) | % missing |
| --- | --- | --- |
| CAUSTON [69] | Commercial microarray (5) | 19–22 |
| DUDLEY [70] | Custom microarray (4) | 5 |
| GARCIA [71] | Custom microarray (1) | 1 |
| HOLSTEGE [33] | Commercial microarray (1) | 12 |
| INGOLIA [17, 72] | RNA-Seq (6) | 4–10 |
| LIPSON [35] | RNA-Seq (6) | 1 |
| LIPSON.MA [35] | Commercial microarray (1) | 4 |
| MACKAY [73] | Custom microarray (1) | 28 |
| MIURA [51] | Competitive PCR (4) | 26–29 |
| NAGALAKSHMI [34] | RNA-Seq (1) | 22 |
| PELECHANO [74] | Custom microarray (1) | 14 |
| ROTH [75] | Commercial microarray (2) | 59–70 |
| VELCULESCU [32] | SAGE (1) | 58 |
| YASSOUR [40] | RNA-Seq (4) | 5 |
| FUTCHER [19] | 2D gel (1) | 99 |
| GHAEMMAGHAMI [27] | Western blot (1) | 34 |
| DEGODOY [37] | LC MS/MS (1) | 25 |
| GYGI [5] | 2D gel (1) | 98 |
| LEE [38] | LC MS/MS (3) | 67–76 |
| LU [16] | LC MS/MS (1) | 83 |
| NAGARAJ [76] | LC MS/MS (6) | 31 |
| NEWMAN [36] | GFP/flow cytometry (1) | 60 |
| PENG [68] | LC MS/MS (1) | 74 |
| THAKUR [77] | LC MS/MS (3) | 84–85 |
| WASHBURN [78] | LC MS/MS (1) | 77 |

Although correlations of replicates within studies are quite high [8], with median *r* = 0.97 for mRNA and 0.93 for protein levels, between-study correlations are far more modest, *r* = 0.62 for mRNA measurements and 0.57 for protein measurements (Figure 1C). That is, data from a typical mRNA study explains 39% of the variance in another study (*r*^2^ = 0.39) and a typical protein study’s results explain only 32% in another study’s variance, consistent with previous studies reporting wide variation between studies [15]. Strong outliers indicate high reproducibility for a two pairs of studies (Figure 1C), but each such outlier is a correlation between separate studies done by the same research group, suggesting the presence of additional variability sources between groups. Coverage of the 5,887 verified protein-coding genes in yeast [39] also varies widely across pairs of studies (Figure 1C).

Coupled with high within-study reproducibility, the low between-study reproducibility indicates the presence of large systematic errors between studies. In a single study [35], mRNA levels in a commercially prepared sample were measured using two methods, a commercial microarray and single-molecule RNA sequencing. These measurements correlate with *r* = 0.86 (73% of the variance explained in one measurement by the other), quite similar to the *r* = 0.84 correlation of the single-molecule measurement with an independent RNA-Seq dataset on RNA from a different study [40]. These data hint, coupled with similar observations in other biological systems [41], that high within-study reproducibility is likely to reflect reproducible biases associated with use of a single measurement technique in addition to reproducible features of the biological sample.

Correlations are modest even between studies using similar methods (e.g., *r* = 0.81 between two RNA-Seq datasets using Illumina instruments [17,40]). Comparing mRNA studies performed using similar or different methods on a shared set of 4,595 genes revealed a consistent bias toward higher median correlations between studies using similar methods, but these differences were not statistically distinguishable (Figure 1D, no *t*-test *P <* 0.05 for differences in correlation when comparing studies employing shared methods versus independent methods after false discovery rate correction).

Between-study correlations quantify the studies’ mean ratio of true variance to total variance, termed the reliability [14,42,43] (see *Methods*). In turn, setting aside sampling error, the maximum observable correlation between any two datasets is equal to the geometric mean of their reliabilities. Because virtually all reported global mRNA-protein correlations involve mRNA and protein levels measured in separate studies, between-study reliabilities are the relevant quantity. The modest reliability values—setting aside those of the same group reporting two studies, which we exclude from this analysis—sharply limit the maximum observable mRNA-protein correlations. This limit has startling consequences: if steady-state mRNA and protein levels actually correlated perfectly (true *r* = 1.0), then given the median observed between-study correlations in Figure 1C, we would expect to observe mRNA-protein correlations of only 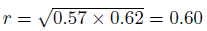.

The data reveal a wide range of modest mRNA-protein correlations with a median of *r* = 0.54 (Figure 2A) quantified either by the Pearson correlation between log-transformed measurements or the nonparametric Spearman rank correlation (Figure S1; both measures produce similar results and we employ the former throughout). The largest pair of datasets covers 4,367 genes and shows an mRNA-protein correlation of *r* = 0.618 (*r*^2^ = 0.38, 38% of protein-level variance explained by mRNA levels), close to consensus values [8]. The largest dataset containing replicated measurements of mRNA and protein in at least two studies yields similar correlation values; notably, averaging paired measurements together and correlating the averages increases the apparent correlation (Figure 2B).

**Figure 2.**
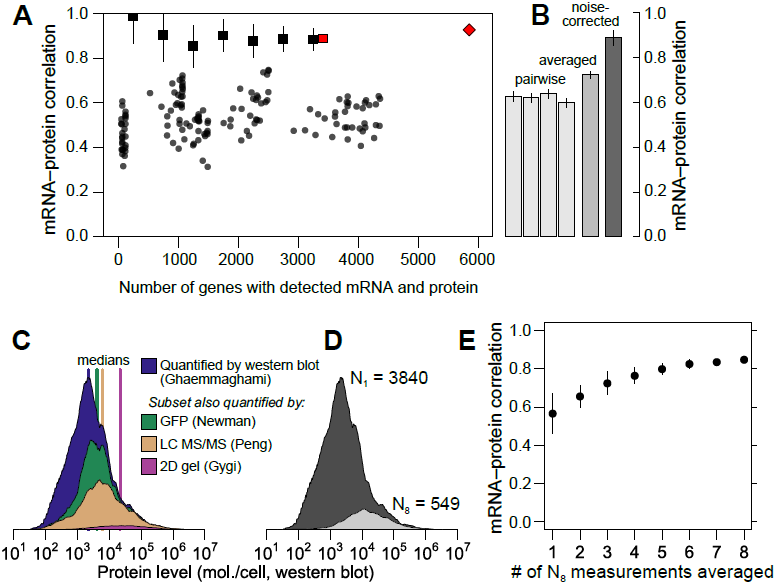
Correlations between mRNA and protein levels vary widely and are systematically reduced by experimental noise. **A**, Datasets vary widely in coverage of 5,887 yeast coding sequences and in resulting estimates of the mRNA-protein correlation. Shown are all pairwise correlations between 14 mRNA and 11 protein datasets, with within-study replicates averaged if present. Correlations are shown between mRNA and protein levels reported without correction (dots); using Spearman’s correction on pairs of datasets (binned, boxes show mean and bars indicate standard deviation); using Spearman’s correction on the largest set of paired measurements (red box); and as estimated by structured covariance modeling for 5,854 genes with a detected mRNA or protein (red diamond). **B**, Correlations obtained for the largest set of paired measurements, two of mRNA and two of protein levels (N=3,418), individually, averaged, and corrected for noise using Spearman’s correction. **C**, Data are missing non-randomly. The distribution of protein levels, in molecules per cell, detected by western blotting [27] are shown, along with the subsets of these data corresponding to proteins detected by GFP-tagging and flow cytometry [36], LC MS/MS [68], and 2D gel [5]. **D**, Distribution of protein-level measurements, assessed by western blotting [27], with at least one protein-level measurement (dark gray, number of genes *N*_1_=3840) and in the subset of genes with at least 8 mRNA and 8 protein measurements (light gray, number of genes *N*_8_=549). **E**, mRNA-protein correlations between averaged mRNA and protein levels over subsets of at most 1, 2, 3, ..., 8 measurements each of mRNA and protein levels drawn at random from the *N*_8_ set. Error bars show the standard deviation of correlations from 50 random samples of the indicated number of measurements.

This averaging effect has a simple explanation: if experimental noise drives down the mRNA-protein correlation, and noise is to some extent random between studies, then averaging together measurements from different studies will increase the correlation as random noise dilutes out and signal titrates in. However, exploiting averaging comes with hidden dangers when using these data. Averaging requires multiple measurements. Few protein datasets cover even half the genome, and incomplete data tend to be biased toward abundant proteins, as revealed by examining levels in a large dataset when restricted to proteins detected in smaller datasets (Figure 2C); it is plausible that higher-expression proteins correlate more strongly with mRNA levels. We therefore checked for an averaging effect using a subset of the data with a minimum level of reproducibility, at least eight mRNA and eight protein measurements, which includes 549 genes. This high-coverage gene subset does encode more highly abundant proteins relative to the rest of the genome as assessed by western blotting (Figure 2D). As a benefit, however, changes in correlation due to averaging within this subset do not merely reflect underlying systematic changes in the expression levels of the analyzed genes. In this subset, the observed mRNA-protein correlation rises markedly as more measurements are averaged together (Figure 2E), more than doubling in the apparent protein-level variance explained by mRNA level (from 33% to 72%) simply by averaging together more measurements of the same genes. These data strongly indicate that experimental noise substantially reduces the apparent correlation between mRNA and protein levels.

### Corrections for noise yield sharply higher correlation estimates

The foregoing analyses involve estimates uncorrected for noise, which as described in the Introduction do not properly estimate the true correlation between the variables being measured. We will first incorporate noise-aware estimates of the true correlation, and then address the more challenging problem of accounting for missing data to arrive at a true genome-scale estimate of the mRNA-protein correlation.

Reduction of correlations by noise can be corrected using information from repeated measurements, assuming the noise is uncorrelated across measurements [22,43]. Quantitative corrections for correlation attenuation were first introduced more than a century ago by Spearman [22], are widely used in the social sciences [43–45], and have found recent applications in biology [14,42,46–49]. Given two measurements each of variables *X* and *Y*, each with uncorrelated errors, the true correlation can be estimated using only correlations between the four measurements *X*_1_, *X*_2_, *Y*_1_, *Y*_2_ (see *Methods*):

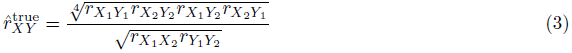

The correction reflects a simple intuition: the denominator quantifies the reliabilities of the measurements, which determine the maximum observable correlation, and the numerator quantifies the observed correlation using a geometric mean of four estimates and is divided by this maximum value to yield an estimate for the true value. The estimate is not itself a correlation coefficient, and may take values outside (–1,1) due to sampling error [43]. Also note that there is no *P-*value associated with this estimate; statistical testing for significant association using uncorrected correlation measures remains valid.

To demonstrate and test Spearman’s correction, we applied it to simulated data generated to mimic key features of mRNA and protein data, but with a known underlying correlation and known measurement reliability. We generated data for 5,000 simulated genes with a range of correlations and fixed reliability; a fixed correlation and a range of reliabilities; and a fixed correlation and reliability with a range of data missing at random, or non-randomly, with a detection bias against low-expression genes. We then measured the observed correlation, uncorrected for noise, and used Spearman’s correction to estimate the true correlation. At each set of parameters, we generated 50 transcriptome/proteome pairs to assess reproducibility.

As shown in Figure 3A–C, noise reduces correlations in a non-negligible way. Given an actual correlation of 0.9, and a reliability of 0.7, higher than the mean values for real data (cf. Figure 1C), the observed correlation has a mean of 0.631 ±0.009 (standard deviation), whereas Spearman’s correction yields a median value of 0.901 ± 0.007, closely matching the true value. Spearman’s correction performs well over a wide range of reliabilities (Figure 3B) and when data are missing at random (Figure 3C), cases where observed correlations provide a wide range of estimates that are all systematically incorrect. Smaller datasets lead to increased variability of the Spearman estimate due to sampling error (Figure 3C). When faced with data biased toward detection of high-abundance mRNAs and proteins, Spearman’s correction systematically underestimates the true correlation (Figure 3D), as expected due to restriction of range effects.

**Figure 3.**
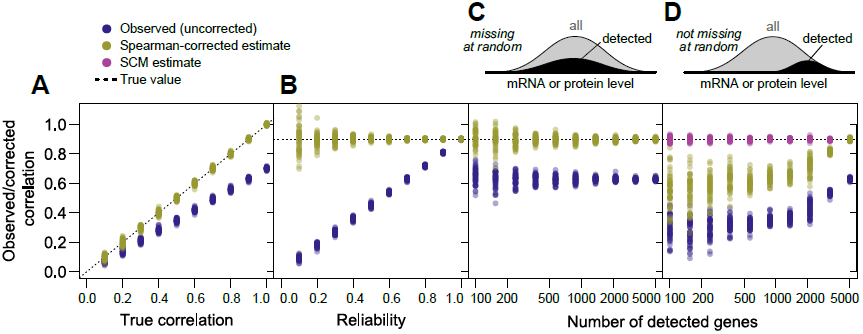
Correlation estimates show widely varying performance on simulated data. (N=5000 “genes”) against the known true correlations used to generate the data (dotted line). 50 replicates were performed at each parameter value. A, Varying true correlation from 0.1 to 1.0 with a fixed reliability (ratio of true to total variance) of 0.7. B, Varying reliability from 0.1 to 1.0 with a fixed true correlation of 0.8. C, Varying the number of detected genes from 100 (2%) to 5000 (100%) with a fixed reliability of 0.7 and fixed correlation of 0.9, with genes missing at random. D, As in C, but with genes missing non-randomly according to the sigmoidal model described in *Methods*, such that low-expression genes are less likely to be detected.

Using Spearman’s correction on real data, we estimated mRNA-protein correlations for pairs of mRNA-and protein-level studies, obtaining a median corrected correlation of 0.92. Variability due to sampling error was large for small datasets as expected (cf. Figure 3C & D), and decreased as dataset size increased, with estimates stabilizing for large datasets (> 3000 genes) at a mean of *r* = 0.88±0.02 (Figure 2A). This value is echoed by consideration of the largest dataset with two mRNA [35,40] and two protein [27, 37] measurements each (Figure 2B). For these data, the four observed mRNA-protein correlations are *r* = 0.60, 0.63, 0.62 and 0.64, and the correlation between mRNA and protein measurements are *r*_mRNA_ = 0.86 and *r*_protein_ = 0.57 respectively, yielding the corrected estimate 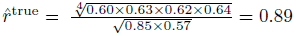.

As demonstrated, Spearman’s correction, while useful, does not address biases due to data that are systematically missing. Spearman’s correction also assumes uncorrelated errors, and thus has no mechanism for handling correlated errors arising due to, for example, protocol similarities within a study or use of similar measurement techniques between studies. Actual datasets show evidence for all of these effects (Figure 1).

### A structured covariance model yields estimates of underlying correlation and of mRNA and protein levels

Extending estimates to the full genome, accounting for structured noise and non-randomly missing data, requires a more sophisticated approach. Even seemingly simple approaches to reduce noise, such as averaging measurements normalized to the same scale, are unworkable as strategies for estimating genome-scale mRNA-protein relationships: only 16 proteins are detected by all 11 protein quantification studies, and these proteins are all highly abundant. Throwing out smaller datasets discards potentially valuable measurements, and it is unclear when to stop, since all datasets are incomplete to some degree.

To address these challenges, we adapted structural equation modeling to admit nonrandomly missing data (see *Methods*). We introduce a structured covariance model (SCM), adapted with important modifications from recent work [25], that explicitly accounts for structured noise arising from replicates and use of shared measurement techniques, explicitly estimates noise at multiple levels and the nonlinear scaling factors linking underlying variables, and allows inferences of latent covariance relationships with imputation of missing data (Figure 4).

**Figure 4.**
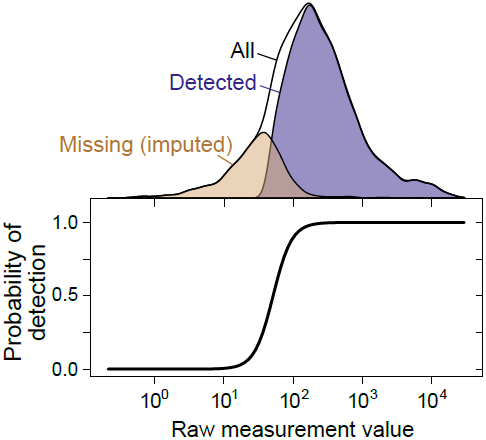
Imputation of non-randomly missing data. The probability of gene or protein detection is modeled in the SCM as an increasing, step-like (logistic) function of the mRNA or protein level (see Methods). Lower panel shows the inferred probability of detection as a function of the measurement value for a single mRNA dataset [69]; top shows the distribution of detected, missing (imputed), and all genes.

The SCM accurately estimates true correlations in simulated data when substantial data are missing nonrandomly, a case on which Spearman’s correction produces severely biased estimates (Figure 3D). Fitting the SCM to real data yields estimates of whole-genome steady-state mRNA-protein correlation of *r* = 0.926 ± 0.004 across all 5,854 genes for which an mRNA has been detected in at least one of the 38 mRNA quantitation experiments (Figure 2A). That is, mRNA levels explain 86% of variation in protein levels at the whole-genome scale. We emphasize that the SCM does not involve any attempt to maximize the mRNA-protein correlation or any assumptions about the strength of the correlation.

To examine the influence of low-coverage datasets on the correlation estimate, we re-fit the SCM on data restricted to studies with no more than 60% or 80% missing values (cf. Table 1), resulting in essentially unchanged correlation estimates of *r* = 0.919 and *r* = 0.933, respectively. Including these smaller datasets does not alter these estimates significantly.

### Comparisons indicate accurate estimates and plausible imputations of mRNA and protein levels

The SCM integrates all data to produces mean and variability estimates of mRNA and protein levels, yielding a dataset in which mRNA levels have been quantified for 5,854 genes and protein levels have been quantified for 4,990 genes in at least one study.

To evaluate the accuracy of these estimates, we linearly scaled them to molecules per haploid cell using high-quality published values for mRNA per cell and protein per cell. Estimates of the number of mRNA molecules per cell range from 15,000 to 60,000 molecules per cell [33,50]. A more recent study argued that the earlier, lower estimate resulted from misestimation of mRNA mass per cell and average mRNA length, with 36,000 molecules per cell as a revised estimate also supported by independent measurements [51]. The higher estimate resulted from rescaling the lower estimate to match expression of five genes measured by single-molecule fluorescence *in situ* hybridization (FISH) [50]. We adopted the 36,169 mRNA molecules per cell estimate [51], and 4*μ*g of protein in 1.5 × 10^6^ cells (2.7pg protein per yeast cell) [52]. Scaled to the latter estimate, SCM protein levels sum to just over 35 million protein molecules per haploid cell, similar to the 50 million molecules per cell estimated previously [19] within the variation in total protein extraction from haploid yeast cells (cf. [53], which estimates 4.95pg per cell).

Scaled SCM per-gene means provide the best point-estimates of molecules per cell (Figure 5A), although the correlation between estimates of means is necessarily higher than the estimated true correlation, since each estimate contains error. For a more representative global view of mRNA and protein levels, we draw a sample from the SCM estimates according to each gene’s mean and variance in levels (Figure 5B). Correlations between sampled mRNA and sampled protein levels (*r* = 0.923) are consistent with the inferred underlying correlation.

**Figure 5.**
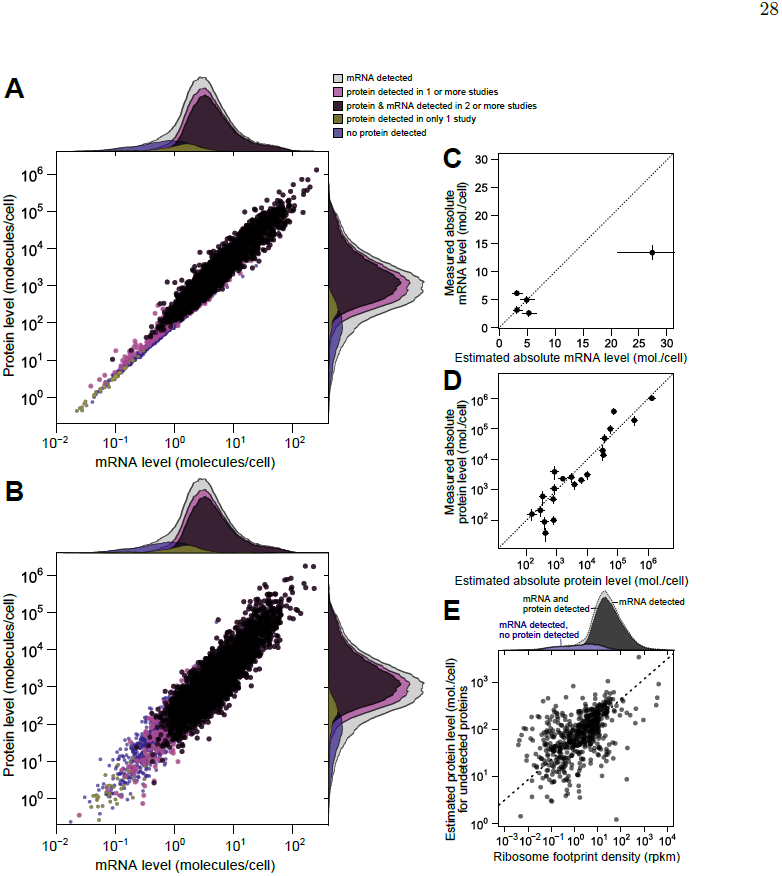
Integrated estimates of mRNA and protein levels using a structured covariance modeling (SCM). **A**, Integrated estimates of mean steady-state protein and mRNA levels across 58 global measurements reveal a strong genome-wide dependence between (r = 0.93). Estimates are produced for any gene with a detected mRNA (gray marginal densities), and other densities characterize subsets by mRNA and protein detection. **B**, A single sample from the SCM estimates provides a representative view of mRNA and protein levels. Colors and marginal densities are the same as in **A. C**, Absolute mRNA level estimates versus single-molecule fluorescence *in situ* hybridization counts [50]. **D**, Absolute protein level estimates versus stable-isotope-standardized single reaction monitoring measurements [54]. Dotted lines in **B** and **C** show perfect agreement. **E**, Evidence for active translation of undetected proteins inferred from ribosome profiling without translational inhibitors [56]. Dashed line shows ranged major-axis regression best fit. Marginal densities show ribosome density for all detected mRNAs (light gray), all mRNAs with a detected mRNA and protein (dark gray), and transcripts with no detected protein (blue). rpkm, reads per kilobase per million reads.

We then compared scaled SCM estimates to small-scale gold-standard, independent measurements of absolute mRNA and protein levels not used in our analysis. (No genome-scale gold-standard measurements of mRNA or protein levels exist for yeast or any other organism.) SCM estimates of absolute mRNA levels matched FISH measurements well [50] (average difference of 1.2-fold between estimated and measured levels [Figure 5B], with one outlier estimate overshooting the FISH value by 1.7-fold). Notably, these results demonstrate that the FISH estimates are compatible with roughly 36,000 mRNA molecules per cell during exponential growth as reported [51], and do not require the almost two-fold higher number of cellular mRNAs extrapolated in the FISH study.

Absolute protein levels for a set of 21 proteins differing up to 25,000-fold in cellular abundance have been measured using single-reaction monitoring (SRM) spiked with stable-isotope standards [54]. SCM estimates correlate better with these absolute levels (*r* = 0.94 between log-transformed values) than does any individual dataset. This includes the only study, which used western blotting [27], which reports levels for all 21 proteins (*r* = 0.90) (Figure 5C, average difference of 1.4-fold between SCM estimates and SRM measurements, compared to 1.8-fold using western blotting). Relative protein levels estimated by integrating multiple datasets using an alternative approach in which noise is not modeled [15] correlate with absolute levels less well (*r* = 0.88) than do the SCM estimates. The structured covariance modeling approach thus estimates steady-state cellular mRNA and protein levels with an unmatched combination of completeness and accuracy.

To evaluate imputation of missing data, we focused on the 864 genes with a detected mRNA but no protein detected in any of the 11 studies. Some of these genes encode well-studied proteins such as the proteasomal regulator Rpn4p and the cyclin Cln3p, indicating clear false negatives. For a systematic evaluation, we turned to ribosome profiling studies [17], which quantify ribosome-protected mRNA (ribosome footprints), providing an estimate of the mRNAs being actively translated *in vivo.* At least one of three studies under compatible experimental conditions [17,55,56] detects ribosomes in the coding sequence of 594 of these 864 genes, suggesting active translation. Normalized ribosome footprint counts for this restricted set of genes correlates with the imputed protein levels (Figure 5E, *r* = 0.51), despite the attenuating effect of range restriction. The most recent study [56] raised concerns about the use of translational inhibitors during sample preparation for ribosome profiling, and we show data collected without inhibitors in Figure 5E. Because the missing protein data correspond to genes at the detection limit of these ribosome-profiling studies, we predict that many of the remaining genes will be found to produce proteins at low levels during exponential growth. The SCM estimates serve as predictions for the levels of these as-yet undetected proteins.

### Translational regulation widens the dynamic range of protein expression

Our results indicate that the true correlation between steady-state mRNA and protein levels in exponentially growing budding yeast is far higher than previously recognized, explaining the vast majority of variation in protein levels on a log scale. In many previous analyses, this would be equivalent to demonstrating a minor role for other forms of regulation: if the variation in protein levels were a pie, and mRNA levels took a slice, other forms of variation would get only the leftovers. As we will show, such competition is largely illusory.

Positive evidence exists for strong post-transcriptional contributions to protein levels. The dynamic range of protein abundance is much wider than mRNA abundance, which must reflect dynamic-range amplification by post-transcriptional regulation [8]. Indeed, wide per-gene variation exists in measurements of translational efficiency [17,55,56].

The report that translational activity, estimated by ribosome profiling, explained more than twice the protein-level variation than did measured mRNA levels [17] prompted us to more closely examine these results. We reproduced these comparisons, and found that subsequent ribosome-profiling studies confirmed the strong predictive power of ribosome footprints for originally employed proteins levels from a single study [37] (Figure 6A). We wondered whether these findings might reflect experimental noise that differed between the mRNA and ribosome-footprint measurements in the original study. Correlations using SCM-integrated protein levels are substantially higher for both SCM-estimated mRNA levels and ribosome footprints measured in all studies, consistent with reduction of noise in the SCM estimates (Figure 6A). SCM-estimated mRNA levels predict protein levels better than any of the ribosome profiling studies (Figure 6A), though this may simply reflect remaining noise and systematic bias in the profiling studies. These results suggest that, contrary to previous reports, there is no evidence that measures of translation have higher predictive power for protein levels than do mRNA levels.

**Figure 6.**
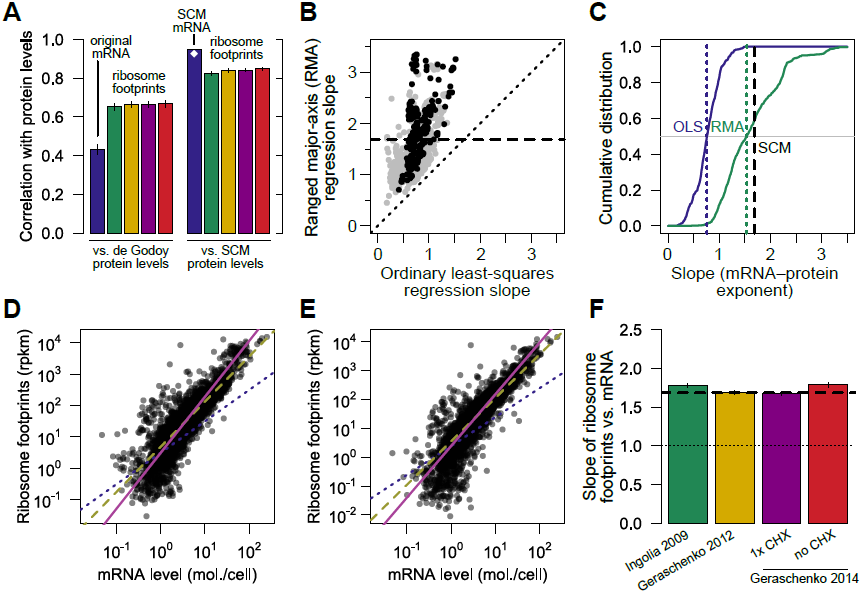
Transcriptional and translational regulation act coherently to set protein levels. A, Left, the correlation of mRNA (blue) and ribosome footprints (green) reported in the original ribosome-profiling study [17] with protein levels used for comparison in that study [37]. Subsequent ribosome footprint datasets ([55], gold; [56], purple and red for measurements with and without cycloheximide, respectively), are shown for comparison. Right, the same comparisons employing SCM-estimated mRNA and protein levels. White diamond shows whole-proteome SCM estimate. Error bars show standard error on the correlation estimate. B, The exponent relating protein and mRNA levels, or equivalently the slope relating log-transformed values, estimated by noise-blind (ordinary least squares, OLS) and noise-aware (ranged major-axis, RMA) regression analyses. Gray points, all pairs of datasets; black points, pairs of datasets covering at least half the detected transcriptome (> 2927 genes). Dotted line shows perfect agreement; dashed line marks integrated SCM estimate (1.69). C, Cumulative distributions of slopes computed by OLS and RMA regression (solid lines), with medians indicated by dotted lines and the SCM slope estimate indicated by a dashed line, cf. Fig. S2. D, Normalized ribosome footprint counts (rpkm, reads per kilobase per million reads, a measure of translational activity) correlates strongly and nonlinearly with mRNA level. Dotted blue line shows linear (slope = 1) fit. Solid magenta line shows RMA regression fit. All regression fits were carried out on the subset of genes with ≥ 1 rpkm. **E**, Same as **D**, using ribosome footprints generated without cycloheximide [56]. **F**, mRNA-ribosome-footprint slopes estimated from independent studies [17,55,56] with and without cycloheximide (CHX).

However, major contributions to protein levels from mechanisms other than mRNA level become obvious upon inspection of the data. The dynamic range of protein expression (from fewer than 50 to more than 1,000,000 molecules per cell [27,54]) is much wider than that of mRNA levels (*e.g.* from 0.1 to 89 molecules per cell in a landmark early study [33]). In the SCM estimates, the range of mRNA expression is roughly 10,000-fold (0.02 to 253 molecules per cell on average), whereas the range of protein expression is more than 1,000,000-fold (an average of 0.4 molecules to 1.3 million molecules per cell). Since both mRNA and protein are roughly lognormally distributed, the ratio of log-transformed ranges, 1.6, yields a rough measure of relative variation. We address more representative estimates of relative dynamic range below. As previously noted [8], this dynamic-range amplification must involve post-transcriptional regulation.

The standard use of a logarithmic scale raises some questions about the interpretation of dynamic range. What does a ten-fold difference mean, if it is between 0.01 to 0.1 molecules per cell rather than between 1 and 10 molecules per cell? Fractional molecules per cell in a population average may indicate mRNAs or proteins present in only a fraction of cells in the population, which can arise in many ways, from conditional expression (*e.g.* during a segment of the cell cycle) to incomplete repression (leakiness). Here, estimates of levels reflect the measurements but confer no particular interpretation. We note that no obvious break or cutoff exists in the data or the SCM estimates to suggest a gene-expression threshold below which the biology changes qualitatively.

### Translational regulation multiplies transcriptional signals with high fidelity

A consequence of two facts—the much higher dynamic range of protein levels than mRNA levels, and the strong log-log linear correlation between the two—is that steady-state protein levels cannot be (even noisily) proportional to steady-state mRNA levels at the genome scale. In the standard model (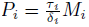 with protein *P* and mRNA *M* for gene *i*, cf. Equation 2), steady-state protein levels will be roughly proportional to steady-state mRNA levels on a log-log scale assuming translation rates and degradation rates are uncorrelated with mRNA levels. This is most easily seen considering the case of constant translation and degradation rates (*τ*_*i*_ = *τ* and *δ*_*i*_ = *δ*, respectively) across all genes, such that 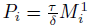 where we have made explicit the exponent of 1. In this case, 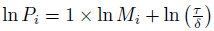. Deviations from proportionality can be captured by deviations from a log-log slope of 1.

As described in the Introduction, several studies have estimated slopes very near 1, but have not accounted for error-induced systematic underestimation of slopes due to regression-dilution bias [28]. We therefore used a noise-tolerant regression technique closely related to principal component analysis known as ranged major-axis (RMA) regression [30], which yielded a range of slopes systematically higher than the ordinary least-squares regression slopes (Figure 6B,(C) and have a median of 1.54. RMA regression is symmetric, such that regression of *Y* on *X* produces the same slope as regression of *X* on *Y.* Other widely used techniques with the same symmetry property but different technical assumptions each yield slopes substantially larger than 1 (Figure S2). The estimated slopes for individual pairs of datasets span a wide range, even using RMA and limiting attention to large datasets (Figure 6B), suggesting the existence of systematic biases, toward increased and decreased variance, separating these studies. The presence of such biases in protein-quantitation studies, though not their precise source, has been previously noted [57].

The SCM approach, which accounts for both noise and missing data, yields an estimated slope of 1.69, compatible with the range of estimates from noise-aware methods on individual pairs of datasets (Figure 6B,C) and also similar to the expectation (1.6) derived from examination of the relative dynamic ranges above. Steady-state protein levels therefore reflect a dramatic multiplication of the transcriptional signal: rather than competing with transcriptional regulation as often assumed, post-transcriptional regulation cooperates.

If translational activity drives much of this cooperative amplification, it must rise nonlinearly with mRNA level, which we assessed by plotting a proxy, ribosome occupancy on mRNA measured by ribosome profiling, against SCM-estimated mRNA levels. A steeper-than-linear relationship is visually clear from examination of the linear fit (slope = 1) compared to the RMA regression line (slope = 1.69, Figure 6D). Data from a separate ribosome-profiling study carried out without the translational inhibitor cycloheximide (CHX) show similar nonlinear behavior (slope = 1.79) (Figure 6E). The results of four studies by two groups yield log-log slopes close to, or above, the SCM estimate, even after restricting slope estimation to ribosome footprint values > 1 to reduce biases possibly introduced by low-count genes (Figure 6E). In summary, measured variation in translational activity correlates strongly with mRNA level and is sufficient to quantitatively account for the nonlinear relationship between mRNA levels and protein levels.

### A toy model illustrates non-independent contributions from transcription and translation

The analysis above illustrates a fundamental asymmetry: although absence of post-transcriptional regulatory processes would produce a perfect mRNA-protein correlation [1], a perfect mRNA-protein correlation need not indicate a negligible post-transcriptional contribution to relative protein levels.

Contrary to the conclusions of many analyses, it is possible for mRNA levels and (for example) translation rates to each explain more than 50% of protein-level variation. Both processes could each contribute 100% of protein-level variation. All that is required is that their contributions not be independent.

To see this, consider the following toy model for regulation of protein levels which does not involve assuming that translation rates are independent of mRNA levels:

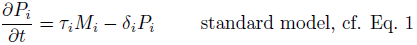

with

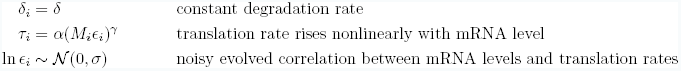

Despite appearances, the functional relationship between translation rates and mRNA levels does not imply or depend on mechanistic properties of transcription and translation. All variance in this model (as in all analyses in the present work) derives from differences between genes, so the functional relationship merely describes an empirical correlation. As described in more detail in the Discussion, such a correlation can arise if genes have evolved differential translational efficiencies tuned to multiply transcriptional signals.

In this toy model, with *ϵ*_*i*_ = 1 (or more generally *σ* = 0), translation rates and mRNA levels reinforce each other perfectly albeit nonlinearly. Under these conditions, steady-state mRNA levels explain 100% of the steady-state protein-level variation on a log scale. Translational regulation also explains 100% of the protein-level variation.

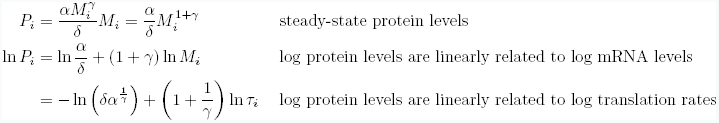

Adding variation to translation rates (*σ* > 0) and fixing other parameters allows close reproduction of the SCM estimates on several dimensions (Figure 7A,B; source code including parameters presented in *Methods*). Both datasets have similar mRNA-protein correlation (*r* = 0.923 vs. *r* = 0.932), similar log-log slopes (1.68 for both), and similar dynamic ranges for mRNA and protein levels.

**Figure 7.**
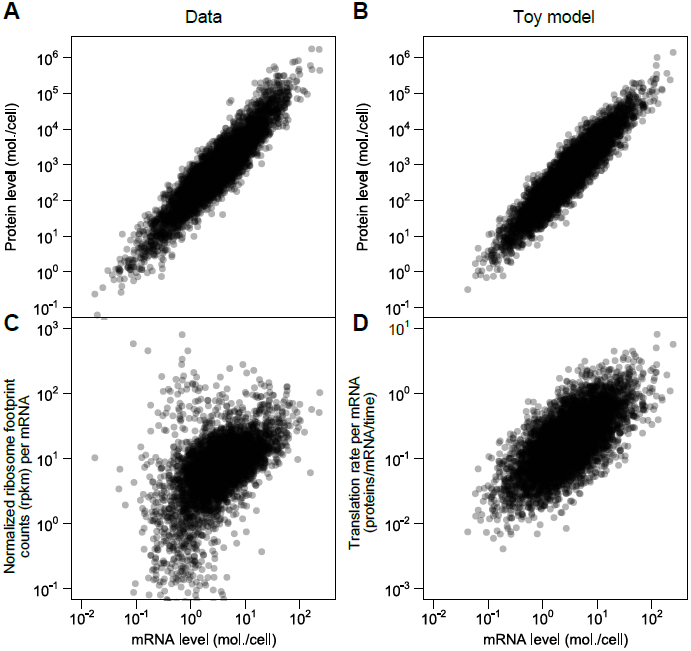
A simplified model captures major features of the steady-state mRNA-protein relationship. A, mRNA and protein levels drawn from the SCM estimates (cf. Figure 5B). B, mRNA and protein levels generated according to a toy model (see text and Methods). C, Normalized ribosome footprint counts (rpkm), averaged across six datasets, per mRNA, compared to steady-state mRNA levels. D, Translation rate per mRNA versus mRNA level in the toy model.

The critical difference between this model and the standard model for protein-level variation, Eq. 1, is the evolved strong positive correlation between mRNA levels and translational activity per mRNA. This, too, is evident in the data, when calculating translational activity per mRNA (frequently called the translational efficiency [17,55]) by dividing ribosome-profiling footprint counts by SCM-estimated mRNA levels (Figure 7C, RMA slope = 0.82, Pearson *r* = 0.53). The correlation is mirrored by the toy model, where translation rate per mRNA and mRNA level can be directly compared (Figure 7D; RMA slope = 0.87, Pearson r = 0.68).

## Discussion

Our results demonstrate that the frequently reported result that steady-state mRNA levels explain less than half (30–50%) of the variation in protein levels constitutes a significant underestimate. In exponentially growing budding yeast, the best-studied system and source of many of these claims, we find that the true value at the whole-genome scale, taking into account the reductions in correlation due to experimental noise and missing data, is closer to 85%.

Many thoughtful studies have tackled this problem before, arriving at results that match ours on certain dimensions, but via quite different approaches. Previous work has employed versions of Spearman’s correction [14], contended with differences in dynamic range by adopting nonparametric approaches [1,16], and integrated multiple datasets [7,10,15,16]. All of these works have reached conclusions which differ from the portrait assembled here.

Our analysis transcends these studies on several fronts. The present study incorporates more measurements than any previous work. We distinguish between correlations between measurements and estimates of underlying correlations accounting for between-study reliability, a critical difference that has largely eluded previous work. The structured covariance model natively handles nonrandomly missing data to provide more complete and accurate molecules-per-cell estimates than previous studies. Most importantly, we have not relied on the common but mistaken assumption that different modes of regulation act independently.

A consistent approach in the literature has been to pit transcriptional and post-transcriptional variation against each other, both analytically and rhetorically (*e.g*., “transcriptional regulation is only half the story” [26]). As we have shown, the data do not fit this competitive paradigm, and even invalidate some of its analytical assumptions, such as independence and non-collinearity. The competitive versus cooperative aspects of post-transcriptional regulation come to the fore when considering the dynamic ranges of gene expression. A wider range of protein than mRNA levels is well-established in a range of organisms [2,14,58], and our results further cement this observation. However, dynamic-range variation could be achieved in different ways, captured by two extremes. At one extreme, post-transcriptional regulatory variation is uncorrelated with transcriptional regulation, reducing the contribution of mRNA levels to protein levels. At the other extreme, post-transcriptional variation correlates strongly with transcriptional regulation, multiplying the transcriptional signal with little interference. In both cases, post-transcriptional regulation amplifies the dynamic range of gene expression, but only in the latter case does it also faithfully amplify the mRNA signal itself. Our data clearly and convergently indicate that the biology, at least for this organism under these conditions, lies toward the latter, cooperative extreme.

Coordinated transcriptional and translational signal amplification may explain a range of other observations, particularly regarding proteins-per-mRNA (PPM) ratios, which are frequently used to isolate signs of post-transcriptional regulation. Because post-transcriptional amplification correlates strongly with mRNA levels, PPM will remain correlated with mRNA, and as a consequence, any sequence features correlated with mRNA will tend to correlate with PPM as well. As an example, amino-acid composition correlates with PPM in yeast [16], with valine/alanine/glycine frequencies higher in high-PPM sequences and leucine/asparagine/serine frequencies lower in high-PPM sequences. These are precisely the same amino acids previously shown to vary most strongly in frequency, in the same directions, with increasing mRNA abundance [59]. Similarly, many other correlates of PPM are also correlates of mRNA levels (codon bias, tRNA adaptation), including mRNA level itself [1,10]. For features such as codon bias, which arises in response to selection for translational efficiency [60], association with increased PPM might seem an obvious causal link, but because codon bias strongly associates with mRNA level, the null expectation is that it will correlate with PPM even if codon bias had no effect on translational activity at all. Analyses of the determinants of protein levels must contend with the collinearity and non-independence of contributing processes.

The strong correlation between steady-state mRNA and protein levels may seem to validate the use of mRNA levels as relatively faithful proxies of protein levels. We urge caution, as a tempting conclusion—that mRNA changes serve as faithful proxies for protein changes—does not follow. Attempts to infer the correlation between mRNA and protein changes from steady-state mRNA-protein correlations confuse two distinct and complex phenomena. The genome-scale relationship between mRNA levels and protein levels is an evolved property of the organism, reflecting tuning by natural selection of each gene’s transcriptional and post-transcriptional controls, rather than a mechanistic input-output relationship between mRNA and protein mediated by the translational apparatus. Two genes with steady-state mRNA levels differing by 10-fold may have 500-fold differences in protein levels due to evolved differences in their post-transcriptional regulation. These evolved steady-state differences do not predict how the protein levels for these genes will change if both mRNAs are induced 10-fold, because evolution does not occur on this timescale; the changes in protein levels are instead dictated by the cellular mechanisms of translation.

An important intermediate case between the evolutionary and mechanistic cases is variation in mRNA and protein levels in individuals across a genetically diverse population. The potential for correlations between mRNA and protein relies upon substantial true variance in mRNA levels. In population-variation studies, one expects relatively few variants and resulting variation far lower than the orders of magnitude considered here. Correspondingly, in such studies mRNA-change-protein-change correlations may be low even given a strong underlying link between mRNA and protein levels.

If the nonlinear multiplication of mRNA levels into protein levels is an evolved property, what mechanism(s) has evolution exploited? The present work supports a particular class: the increased density of ribosomes on high-expression mRNAs, with variation sufficient to account for the nonlinearity, suggests increased rates of translation initiation as the major contributor. Correspondingly, recent work has shown that in yeast and a wide range of other organisms, the stability of mRNA structures in the 5’ region weakens as expression level increases, favoring more efficient translation initiation [61], and wide variation in heterologous protein levels can be achieved by varying mRNA stability near the initiation site [62,63].

Several limitations still attend our approach. By assuming single multiplicative errors per experiment, we ignore variation in per-gene error which may be systematically different between low-and high-expression genes and/or systematically affect particular measurement techniques [57]. For example, limitations in the dynamic range of a measurement technique will tend to compress the resulting measurements, causing such systematic errors. Our model does not contend with distortions possibly imposed by alterations to 3’ regulatory signals (*e.g.* tagging with affinity epitopes [27] or fluorescent proteins [36] to enable protein detection), or with variability in quantification due to propensities of particular mRNAs to be more efficiently sequenced or for their protein products to be unusually amenable to mass-spectrometric detection. The lack of any gold-standard genome-scale measurements hinders detection of such biases. Our results underscore the urgent need for such standard measurements of absolute mRNA and protein levels to enable identification and correction of systematic errors in established and emerging gene-expression measurement techniques. More sophisticated models for experimental error at many levels, which would be informed by but need not wait for such gold-standard measurements, also promise to provide higher-fidelity biological estimates from existing data.

We infer a higher mRNA-protein correlation (*r* = 0.93) here than when using an earlier, related model [25] (*r* = 0.82), a difference we attribute to two factors. First, the present analysis stratifies by measurement technology using all data, whereas the previous estimate did not, although in that study, stratifying by technology on a reduced dataset yielded *r* = 0.86 [25]. Here, using all data and treating technology-related experimental noise separately from other sources of noise, we are able to average out more systematic technology biases, likely producing superior estimates of the associated measurement variability and reducing noise-induced attenuation of the mRNA-protein correlation. Second, in the present analysis, population-averaged protein levels and mRNA levels are constrained to each have a single underlying variance, whereas in the earlier study each experimental replicate had a separate variance. Inference of artificial experiment-specific variances spread variability across experiments (overfitting), where in the present analysis, we adopt the more biologically plausible stance that the true underlying mRNA and protein population-average distributions each have a characteristic variance which is measured by each experimental replicate. The present model, deprived of extra parameters, infers higher correlations.

Our study considers a single well-studied growth condition for a single well-studied organism, raising questions about how to generalize this work. The principles of accounting for noise, but not precise results, can and should be extrapolated to regulatory contributions in other settings and other organisms. An influential study on mouse fibroblasts measured mRNA and protein levels and degradation rates for thousands of genes [2], concluding that mRNA levels explained 41% of the variation in protein levels, with most variation instead explained by translational regulation. Our results indicate many ways in which the results of this study could be profitably revisited. Indeed, a recent follow-up study concluded that, once effects of error and missing data were accounted for, mRNA levels explain 75% or more of the protein-level variation in these data [20].

The protein regulatory environment of rapidly dividing cells differs from that of many other cellular states. The faster cells divide, the more rapidly protein molecules partition into daughter cells, adding an approximately constant amount to all protein removal rates and consequently reducing between-gene variation in these rates. This will tend to increase the dependence of protein levels on mRNA levels, and decrease the dependence on degradation rates, during proliferation.

In addition to cellular state, regulatory contributions depend on timescale. Post-transcriptional processes must dominate protein-level changes within seconds to a few minutes of a stimulus or signal; transcriptional responses, particularly in eukaryotes, where transcription and translation are uncoupled, are all but powerless at this timescale. As such, the notion of general determinants of protein levels without regard to timescale has questionable utility. A final theme emerging from our study is that careful empirical studies, coupled with noise-aware analyses, are needed to determine regulatory contributions for any cellular condition of interest at any timescale.

## Methods

### Reliability

Let us assume we wish to measure latent variables *ϕ* and *ψ* but, due to noise, actually observe variables *X* =*ϕ* + *ϵ*_*X*_ and *Y* = *ψ* + *e*_*Y*_ where the random noise variables *ϵ*_*X*_ and *ϵ*_*Y*_ are uncorrelated and mean zero. The reliability

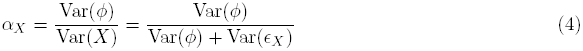

quantifies the ratio of signal variance to total (signal plus noise) variance in *X.* Given two random variables *X*_1_ and *X*_2_ representing replicate measurements of *ϕ*, the latent (true) variance is Cov(*X*_1_, *X*_2_) = Cov(*ϕ* + *ϵ*_*X*1_, *ϕ* + *ϵ*_*X*2_) = Cov(*ϕ,ϕ*) = Var(*ϕ*), where the error terms vanish because they are uncorrelated by assumption. Thus, the Pearson correlation between replicates is

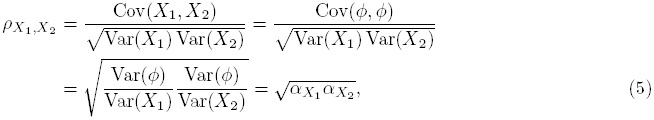

which is the geometric mean of the reliabilities of the two measurements.

### Spearman’s correction

We wish to infer the Pearson correlation coefficient between latent variables 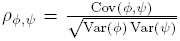 but, due to noise, we observe random variables

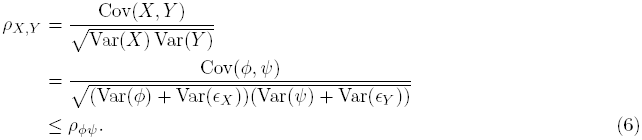

with equality only when Var (*ϵ*_*X*_) = Var(*ϵ*_*Y*_) =0 (*i.e.* there is no noise).

Uncorrelated noise has no average effect on the numerator because errors cancel (see above), but the error terms in the denominator do not cancel. This effect additively inflates the variances in the denominator, biasing the observed correlations downward relative to the truth. Given the reliabilities *α*_*X*_ and *α*_*Y*_, Spearman’s correction is given by

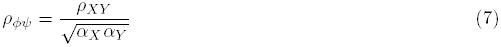

To estimate *ρϕψ*, we need estimates of *ρ*_*XY*_, *α*_*X*_ and *α*_*Y*_. A natural estimator replaces these population quantities with the sample correlation coefficients, *r*_*xy*_ 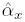 and 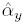 with

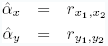

where *x*_1_, *x*_2_ are realizations of *X* and *y*_1_, *y*_2_ are realizations of *Y.* These replicates are used to estimate reliabilities.

The true correlation, *ρ*_ϕ,ψ,_ can then be estimated using only correlations between measurements:

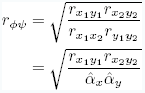

We extend this estimate to

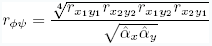

which has the further desirable properties of exploiting all pairwise correlations and being independent of the choice of indices.

### Data collection

We gathered 38 measurements from 13 studies measuring mRNA expression, and 20 measurements from 11 studies measuring protein concentrations, yielding a total of 58 high-throughput measurements of mRNA and protein levels from a maximum of 5,854 genes in budding yeast. The measurements were taken using different technologies including custom and commercial microarrays, competitive PCR, high-throughput RNA sequencing, flow cytometry, western blotting, scintillation counting of ^35^S-labeled protein on 2D gels, and liquid chromatography coupled to tandem mass spectrometry (LC-MS/MS) using a range of labeling and quantification techniques. All yeast cultures were haploid *S.cerevisiae* growing in shaken liquid rich medium with glucose between 22°C and 30°C and sampled during the exponential growth phase. Details of the datasets are summarized in Tablenon 1.

For analytical purposes, we treat data from one study [35] which performed two independent measurements using different methods as two studies (RNA-Seq and microarray), one per method. This study’s RNA-Seq employed a single-molecule sequencing method, smsDGE; we treat this as an RNA-Seq dataset.

Raw data (with missing values), data normalized and imputed using the SCM, and merged moleculesper-cell estimates are archived in Dryad (http://datadryad.org) with DOI doi:10.5061/dryad.d644f.

We downloaded ribosome-profiling data from the primary sources [17,55,56].

### Statistical analysis

All analyses were carried out using R [64] using custom scripts which may be downloaded from GitHub (http://github.com/dad/mrna-prot). Regression analyses using major-axis (MA), scaled major-axis (SMA), and ranged major-axis (RMA) regression were performed using the package lmodel2. RMA was performed using interval ranges.

### The structured covariance model (SCM)

The model has two components: an observation model *p*(*I*_*i,j*_|*X*_*i,j*_), which provides the probability of observing a value for mRNA/protein *i* in replicate *j*, given the underlying mRNA/protein level, and a hierarchical model *p*(*X*_*,j*_|…) for the underlying mRNA/protein levels themselves. The full model is specified as

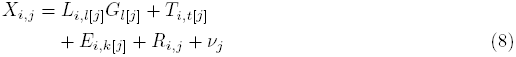

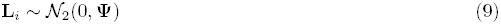

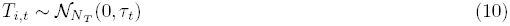

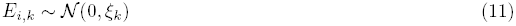

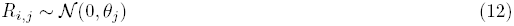

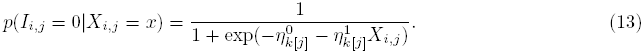

Random variables *L*_*i.l*_ correspond to the true denoised protein (*l* = 1) and mRNA (*l* = 2) levels, for mRNAs and proteins *i* = 1,...,*N*, and L_*i*_ = [*L*_*i,1*_, *L*_*i,2*_]’. The random variables *T*_*i,t*_ and *E*_*i,k*_ capture common technological variation and batch effects, respectively, *t* = 1,..., *N*_*t*_, *k* = 1,..., *N_E_. Ri,j* are experimental noise for replicate *j* = 1,..., *N*_*R*_.

Both technology effects and batch effects between experiments are assumed to be independent, 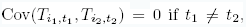 and 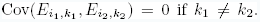 Measurement noise is independent between replicates, 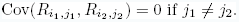

The parameters *v*_*j*_ corresponds to the normalizing constants of the mRNAs/proteins within a replicate (on the log-scale, normalizing constants become offsets). The coefficient *G*_*l*_ represents the log-variance of the denoised true mRNA or protein. The ratio 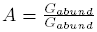 represents the amount of post-transcriptional amplification of mRNA to protein. At steady state we expect

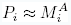

for protein *P*_*i*_ and mRNA *M_i_.*

This model falls into the class of models that were extensively studied in an earlier work [25]. The results are largely insensitive to deviations from parametric modeling assumptions and to several details of prior specifications.

### Missing data model

Equation 13 models the probability that measurement *X*_*i,j*_ is missing, *p*(*I*_*i,j*_ = 0|*Xi,j* = *x*), as a logistic function of the value of the measurement. This data is *not* missing at random (NMAR) since the probability of missingness is a function of the (possibly missing) value. Such a missingness model is said to be *non-ignorable.* The parameters of the missing data mechanism, 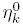 and 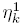, and value, *X*_*i,j*_, uniquely determine the probability that the measurement is observed. For instance, when 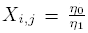, then the missingness probability *p*(*I*_*i,j*_ = 0|*X*_*i,j*_ = *x*) = 0.5.

### Prior specifications

To complete the model specifications we place priors on **Ψ**, *τ*_*t*_, *ξ*_*k*_, *θ*_*j*_, 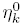 and 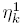. We use either flat or weakly informative priors on all parameters so as to bias the inference as little as possible. For the parameters 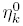 and 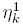 of the logistic observation model we use a Cauchy prior with mean zero and scale 2.5 as suggested by [65]. The role of this prior is to regularize the slope of the logistic regression in cases that have a very sharp cutoff. We assume flat priors on the scaling factors, G*_k_*, and the measurement bias parameters *v_j_.* For the replicate and experiment variances *θ_j_* and *ξ_k_* we use independent conjugate Inv-Gamma(3/2, 3/10) prior. Finally, for the estimand of interest, we assume **Ψ** is a priori drawn from the set of correlation matrices with marginally uniform correlations [66]. The priors for the variances are standard Inverse-Gamma priors, and they are very weak. They correspond to three degrees of freedom, i.e. three data points, and a scale of 1/5. Essentially their role is avoid extreme values at the beginning of the MCMC chain. The prior on the correlation **Ψ** is critical, since this the main quantity of interest. The standard prior for normal covariance matrices is an Inverse-Wishart distribution. For this prior the variances (fixed to 1 in our case) are associated with the correlations. To avoid a strong prior influence on the correlation, we used a prior that is uniform, as suggested by [66].

### Algorithm

MCMC inference in the SCM is done using a Gibbs sampler. The exact conditional draws performed in each time step are:

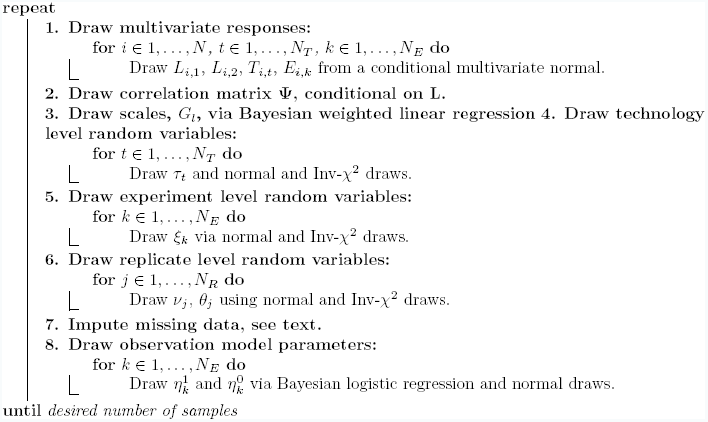

The draws are consistent with standard Bayesian linear regression or logistic (step 8) regression with conjugate variance draws (steps 4–6). [67, Sec. 14.2]. Step 1 is a simple multivariate normal draw, and the imputation in step 7 is done using a Metropolis-Hastings independence sampler. Step 2 is also a Metropolis-Hastings sampler using a random walk proposal, see [66] for the complete method.

### Sensitivity analysis

Our model belongs to a class of models extensively studied by [25], and in the following we summarize the properties and results from these models here.

In simulation studies with data sets generated from the model, the model has good frequentist coverage properties, especially for the *ψ*_1,2_ mRNA-protein correlation parameter.

The model is robust to departures from normality, and the inferred correlation has a very small bias for data sets that are generated from skewed and/or heavy-tailed distributions.

In our model we assume independent observations for different genes and proteins. Genes may have correlated fluctuations, for example if cultures are grown in ways which systematically induce or repress particular pathways. Simulations show that even if a large number of genes are strongly correlated, the inferred correlations are only slightly biased.

The logistic observation model is also robust to mis-specification. In particular, the inferred correlation shows no bias for data generated from a two-stage observation model, with an additional stage in which proteins are missing uniformly with a 0.2 probability.

The model is also minimal, in the sense that the major assumptions of correlated noise and non-ignorable missingness are required to recover the correct mRNA-protein correlation in simulation studies.

### Toy model

Below is R code to reproduce the toy model in Fig. 7.

~~~
# Random number seed
set.seed(115)
# Number of genes
n <-5854
# Exponent of empirical (evolved) relationship between steady-state
# mRNA levels and translation rates
gamma <-0.56
# Scaling factor, 1/time
alpha <-0.1
# Degradation rate, 1/time
delta <-0.001
# Standard deviation of mean-zero variation added to log mRNA levels to yield
# unscaled log translation rates
extra.variation <-1.3
# Steady-state mRNA levels in molecules/cell (log-normal)
# Mean and variance are equal to those of the SCM mean estimates
log.m <-rnorm(n, mean=1.09, sd=1.19)
m <-exp(log.m)
# Translation rate -- add log-normal variation to, and scale, mRNA levels
tau <-alpha*exp(log.m + rnorm(n,mean=0,sd=extra.variation))^gamma
# Steady-state protein levels in molecules/cell (log-normal)
p <-(tau/delta)*m

# Plot protein vs. mRNA
plot(m, p, log=’xy’, pch=16, las=1,
xlab=’mRNA level (mol./cell)’, ylab=’Protein level (mol./cell)’)
# Plot translation rate vs. mRNA
plot(m, tau, log=’xy’ pch=16, las=1,
xlab=’mRNA level (mol./cell)’,
ylab=’Translation rate per mRNA (proteins/sec)’)
~~~

## Acknowledgments

We thank S. Nesterko for early work, N. Ingolia for providing data, and E.W.J. Wallace, K. Cook, and many other colleagues for thoughtful discussions.

---

**Figure S1.**
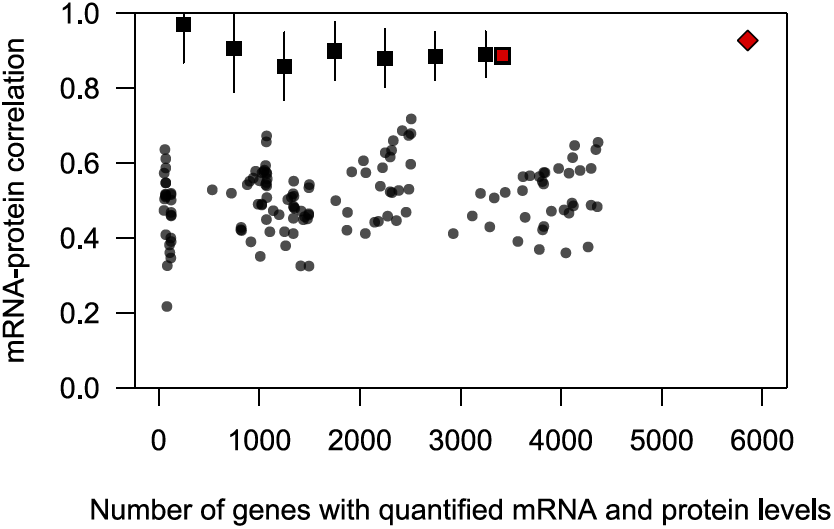
Use of the nonparametric Spearman rank correlation yields similar results to use of the Pearson correlation on log-transformed values (labels as in Fig. 2A). The results of Spearman’s correction on the largest set of paired datasets (red square) and structured covariance model (SCM) fitting (red diamond) are provided for reference.

**Figure S2.**
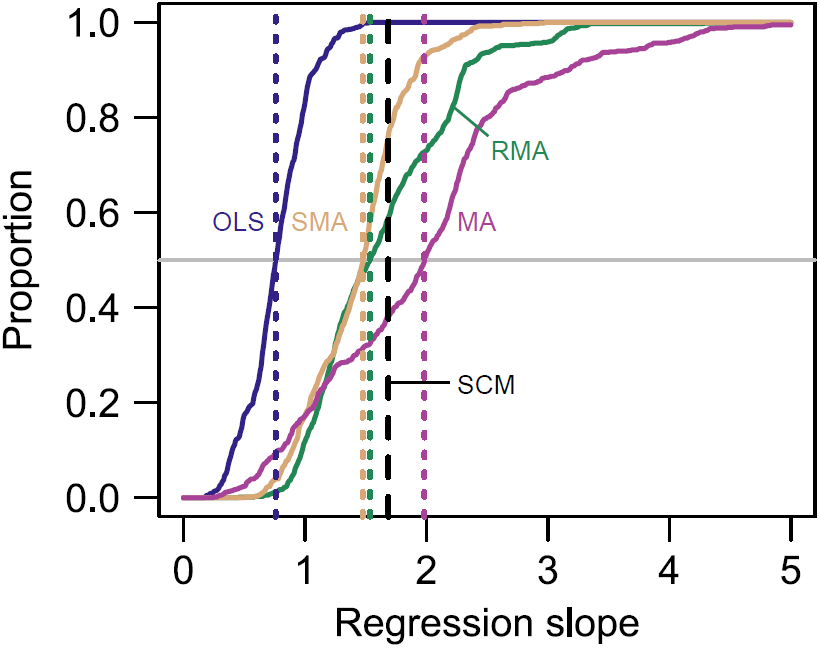
Three methods for slope estimation in the presence of noise yield substantially higher estimates than ordinary least-squares (OLS) regression. Shown are the results of major-axis (MA), scaled major-axis (SMA), and ranged major-axis (RMA) regression of protein levels on log mRNA levels, all values log-transformed but otherwise raw, with slopes extracted and shown as cumulative distributions. The SCM fit value is provided for reference (black dashed line).

